# Persistent Cellular Immunity to SARS-CoV-2 Infection

**DOI:** 10.1101/2020.12.08.416636

**Authors:** Gaëlle Breton, Pilar Mendoza, Thomas Hagglof, Thiago Y. Oliveira, Dennis Schaefer-Babajew, Christian Gaebler, Martina Turroja, Arlene Hurley, Marina Caskey, Michel C. Nussenzweig

**Author notes:** Send correspondence to Gaëlle Breton, Michel Nussenzweig. Equal contribution (Gaëlle Breton, Pilar Mendoza, Thomas Hagglof).

## Abstract

SARS-CoV-2 is responsible for an ongoing pandemic that affected millions of individuals around the globe. To gain further understanding of the immune response in recovered individuals we measured T cell responses in paired samples obtained an average of 1.3 and 6.1 months after infection from 41 individuals. The data indicate that recovered individuals show persistent polyfunctional SARS-CoV-2 antigen specific memory that could contribute to rapid recall responses. In addition, recovered individuals show enduring immune alterations in relative numbers of CD4^+^ and CD8^+^ T cells, expression of activation/exhaustion markers, and cell division.

**Summary:** We show that SARS-CoV-2 infection elicits broadly reactive and highly functional memory T cell responses that persist 6 months after infection. In addition, recovered individuals show enduring immune alterations in CD4^+^ and CD8^+^ T cells compartments.

## Introduction

Individuals infected with SARS-CoV-2 develop both cellular and humoral immune responses to the virus (Robbiani et al., 2020, Gaebler et al., 2020, Suthar et al., 2020, Ni et al., 2020, Grifoni et al., 2020, Peng et al., 2020, Sekine et al., 2020, Rydyznski Moderbacher et al., 2020, Braun et al., 2020, Zhou et al., 2020). Antibody responses include most of the structural proteins expressed by the virus and neutralizing antibodies directed primarily to the receptor binding domain of the spike trimer (S) (Robbiani et al., 2020, Gaebler et al., 2020, Suthar et al., 2020, Ni et al., 2020, Lumley et al., 2020). Cellular immune responses can be broad but vary widely (Grifoni et al., 2020, Peng et al., 2020, Sekine et al., 2020, Rydyznski Moderbacher et al., 2020, Braun et al., 2020, Zhou et al., 2020), and lymphopenia is a prominent feature of more severe infection, affecting CD4^+^ and CD8^+^ T cells, as well as B cells (Tavakolpour et al., 2020, Chen and John Wherry, 2020, Huang et al., 2020, Giamarellos-Bourboulis et al., 2020, Tan et al., 2020, Kuri-Cervantes et al., 2020, Mathew et al., 2020).

Careful studies of immune phenotypes in moderate, severe and recovered individuals revealed T cell responses, ranging from undetectable to robust CD8^+^ T cell and/or CD4^+^ T cell activation and proliferation (Grifoni et al., 2020, Peng et al., 2020, Braun et al., 2020, Neidleman et al., 2020). During acute infection, T cells displayed a highly activated cytotoxic phenotype, whereas convalescent patients harbored polyfunctional SARS-CoV-2-specific T cells that display a stem like memory phenotype (Sekine et al., 2020, Neidleman et al., 2020, Weiskopf et al., 2020). In addition, there is cross-reactivity with seasonal/common cold coronaviruses that suggests that these responses may be associated with a milder clinical course (Sekine et al., 2020, Le Bert et al., 2020).

Coronaviruses elicit variable levels of persistent Immunity to other coronaviruses. For example, individuals infected with MERS remain immune for only 1–3 years, while protection from seasonal coronaviruses is short-lived (Wu et al., 2007, Edridge et al., 2020, Tang et al., 2011). Although there is increasing evidence that cellular immunity plays a major role in resolution of COVID-19, little is known about the persistence of cellular immunity to SARS-CoV-2 (Rodda et al., 2020). This is a particularly important issue when considering an individual’s ability to resist a second exposure to the virus. To determine whether cellular immunity to SARS-CoV-2 persists half a year after infection we examined paired samples obtained an average of 1.3 and 6.1 months after infection from a cohort of 41 COVID-19-convalescent volunteers (Robbiani et al., 2020, Gaebler et al., 2020). All of the individuals tested had RT-PCR confirmed SARS-CoV-2 infection or were close contacts that had seroconverted and were RT-PCR negative at the second time point (Gaebler et al., 2020). The 41 individuals were 63.4% male and 36.6% female, representing a range of different ages, 24-73 years old, and disease severity but skewed to mild forms of the disease (Table 1).

## RESULTS

### Changes in Circulating T cells After COVID-19

The phenotypic landscape of circulating T cells was determined by high-dimensional flow cytometry at both time points and compared to pre-COVID-19 samples from healthy individuals (n=20). Global high-dimensional mapping with t-distributed stochastic neighbor embedding (tSNE) revealed significant persistent alterations in SARS-CoV-2 recovered individuals (Fig. 1A). Relatively under-represented T cell clusters in recovered individuals included Clusters 1, 2, 3, 4, and 14 (Fig. 1B-D and Suppl. Fig. 1B). Clusters 1,3 and 4 showed a phenotype consistent with CD4^+^ central memory T cells (Tcm, CD45RA^-^CD27^+^CCR7^+^, Fig. 1B-C and Fig. S1A). In contrast, cluster 14 displayed features of CD8^+^ Tcm (CD45RA^-^CD27^+^CCR7^+^, Fig. 1B-C and Fig. S1A). Examples of clusters showing a relative increase over control included clusters 10 and 13 (Fig. 1D and Suppl. Fig. 1B). Cluster 10 resembles CD8^+^ T effector cells (Te, CD45RA^+^CD27^-^CCR7^-^, Fig. 1B-D and Fig. S1A) and cluster 13 CD8^+^ T effector memory cells (Tem, CD45RA^+^CD27^-^CCR7^-^, Fig. 1B-D and Fig. S1A). The relative distribution of all of the clusters described above remained abnormal at the 6.1 month time point (Fig. 1D and Suppl. Fig. 1B). We conclude that there are significant shifts in circulating CD4^+^ and CD8^+^ T cell compartments that persist for half a year after SARS-CoV-2 infection.

**Fig. 1.**
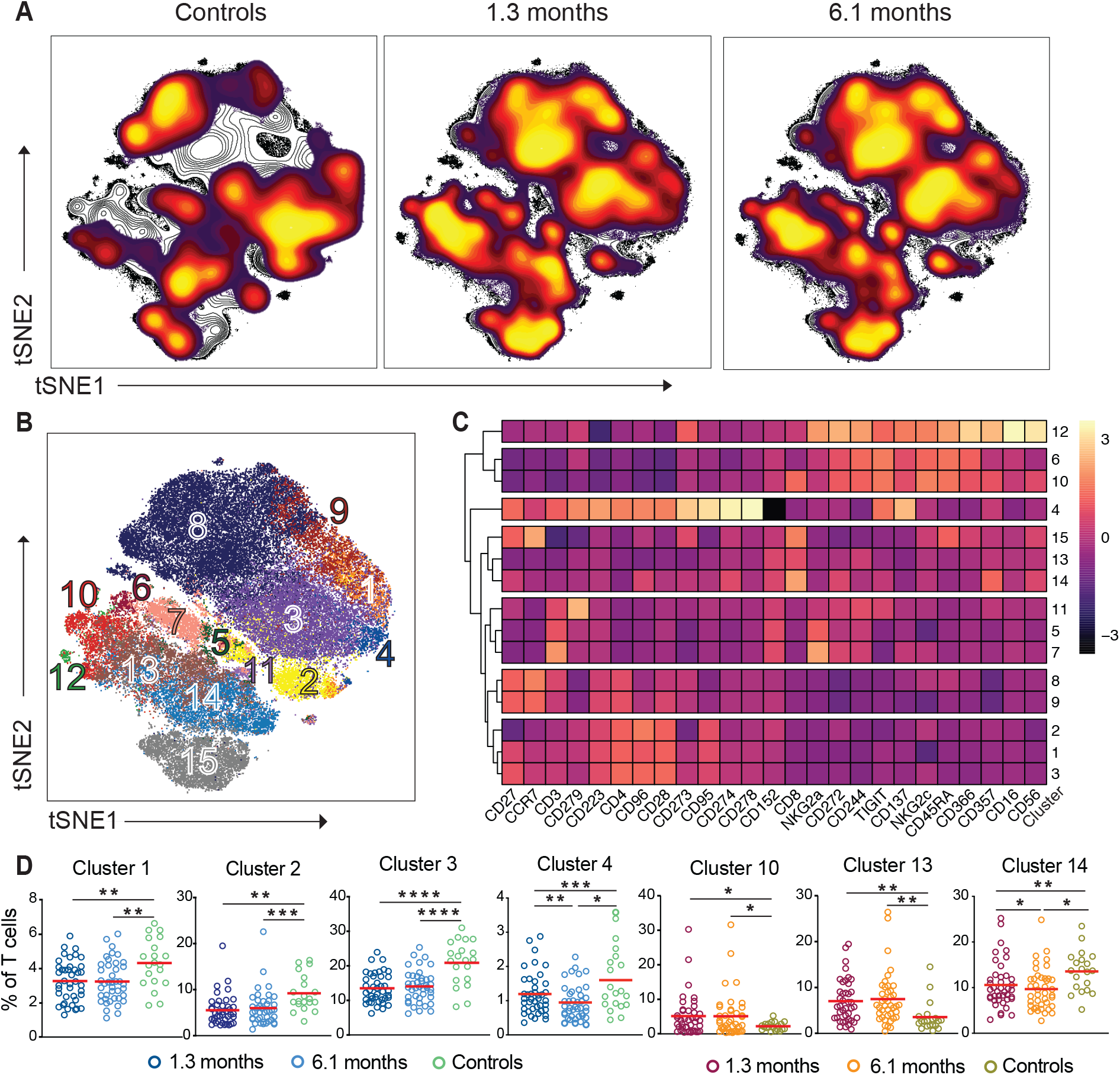
Persistent longitudinal changes in the phenotypic landscape of T cells in individuals recovered from COVID-19. (**A**) Global viSNE projection of pooled T cells for all participants pooled shown in background contour plots, with overlaid projections of concatenated controls, convalescent patients 1.3 months, and convalescent 6.1 months, respectively. (**B**) viSNE projection of pooled T cells for all participants of T cell clusters identified by FlowSOM clustering. (**C**) Column-scaled *z*-scores of median fluorescence intensity (MFI) as indicated by cluster and marker. (**D**) Frequency of T cells from each group in FlowSOM clusters indicated. Each dot represents an individual with COVID-19 at 1.3 months (dark blue for CD4^+^ T cells and dark red for CD8^+^ T cells) or 6.1 months (light blue for CD4^+^ T cells and orange for CD8^+^ T cells) as well as control individuals (green).

To further examine changes in the T cell compartment we queried the data by traditional gating (Suppl. Fig. 2). The relative proportions of circulating CD4^+^ T cells decreased significantly 1.3 months after infection whereas the circulating CD8^+^ T cells increased; but both returned to near physiologic levels by 6.1 months (Fig. 2A and 2B). PD-1 expression is modulated on activated and exhausted T cells and is associated with acute and prolonged changes in T cell function after viral infection in mice (Schonrich and Raftery, 2019, Jubel et al., 2020). PD-1 expression was decreased on CD4^+^ and CD8^+^ T cells at both time points (Fig. 2C and 2D). Consistent with these alterations, TIGIT, Tim-3 and CD25 expression were also abnormal (Fig. 2C and 2D). We conclude that there are persistent changes in the distribution of circulating CD4^+^ and CD8^+^ T cells and their expression of activation/exhaustion markers.

**Fig. 2.**
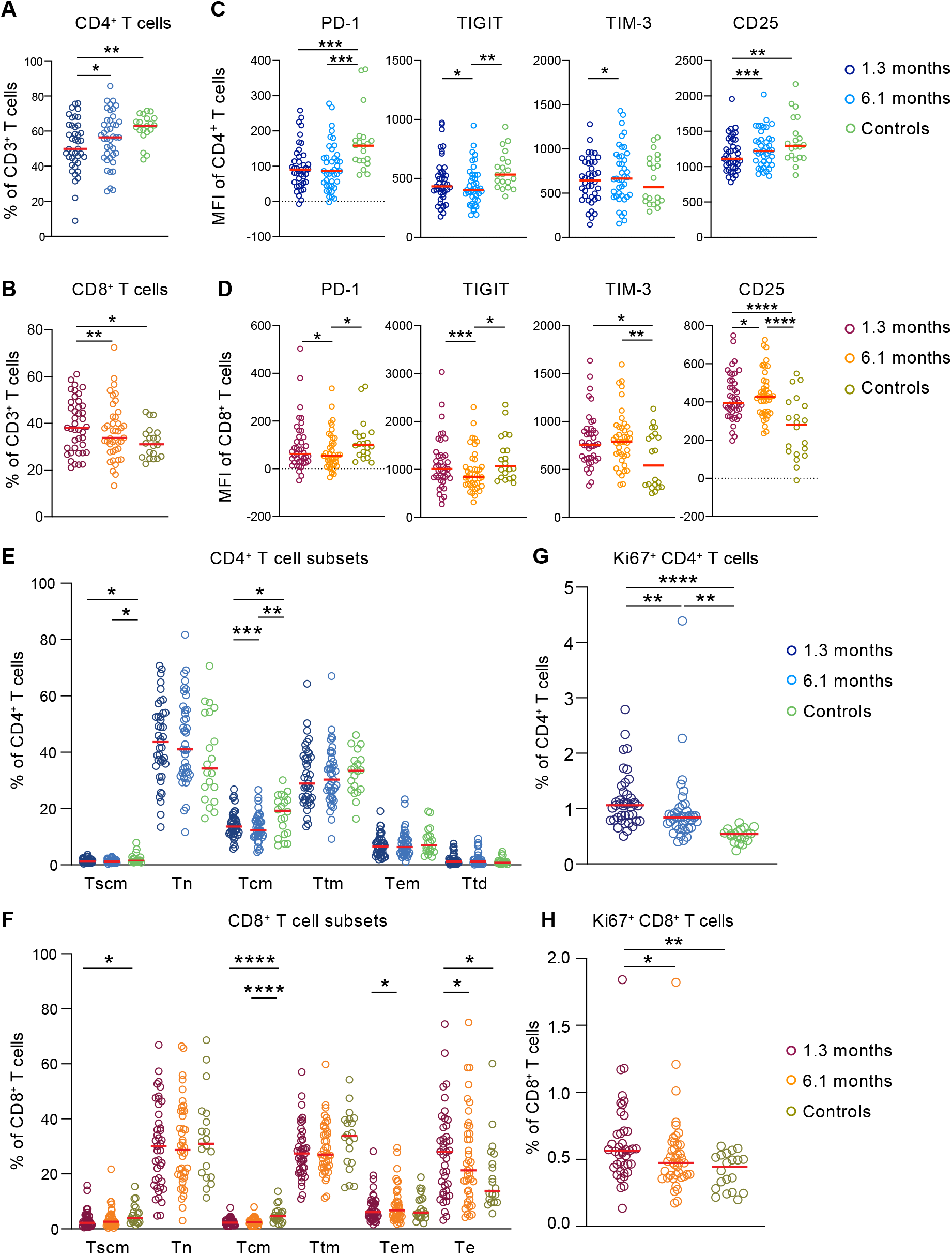
T cell abnormalities persist after 6.1 months. (**A**) Frequency of CD4^+^ T cells out of total CD3^+^ T cells. (**B**) PD-1, TIGIT, TIM-3 and CD25 expression of CD4^+^ T cells. (**C**) Frequency of CD8^+^ T cells out of total CD3^+^ T cells. (**D**) PD-1, TIGIT, TIM-3 and CD25 expression of CD8^+^ T cells. (**E**) Percentage of Tscm, Tn, Tcm, Ttm, Tem, and Ttd CD4^+^ T cells. (**F**) Percentage of Tscm, Tn, Tcm, Ttm, Tem, and Te CD8^+^ T cells. (**G**) Frequency of cycling CD4^+^ T cells. (**H**) Frequency of cycling CD8^+^ T cells. Each dot represents an individual with COVID-19 at 1.3 months (dark blue for CD4^+^ T cells and dark red for CD8^+^ T cells) or 6.1 months (light blue for CD4^+^ T cells and orange for CD8^+^ T cells) as well as control individuals (green). Significance determined by paired t test for comparisons between time points within individuals and unpaired t test for comparison between controls and COVID-19 individuals. *p<0.05, **p<0.01, ***p<0.001, ****p<0.0001.

Consistent with the clustering analysis, central memory CD4^+^ and CD8^+^ T cells decreased, and this defect persisted throughout the observation period (Fig. 2E and 2F). In addition, there was an increase in cycling CD4^+^ and CD8^+^ T cells (Ki67^+^) at both time points (Fig. 2G and 2H). Both central memory and cycling cells also showed lower levels of PD-1 expression at both time points (Suppl. Fig. 3A-D). In contrast, we found no significant changes in circulating T_FH_ or T_reg_ cells (Suppl. Fig. 4). Thus, the more traditional gating strategy is generally consistent with the tSNE analysis.

### SARS-CoV-2-antigen specific CD4^+^ T cells

To investigate SARS-CoV-2-antigen specific CD4^+^ T cells, we stimulated the cells with a collection of pooled SARS-CoV-2 peptides *in vitro*. COVID-19 convalescent individuals were compared to healthy donors by tSNE using high dimensional flow cytometry. Antigen-specific CD4^+^ T cells expressing memory markers as well as IL-2, IFN-γ, TNF-α and CD154 were markedly increased in COVID-19 recovered individuals compared to healthy donors, but the relative frequency of these cells decreased at the 6.1 month time point (clusters 2, 3, 4 and 6, Fig. 3A-D and Suppl. Fig. 5A). The magnitude of the decrease in these responses between the time points varied between 22-32% depending on the cluster (Fig. 3D) and on the individual peptide pool (Suppl. Fig. 5B). In contrast, responses to CMV peptides remained unchanged between the 2 time points (Suppl. Fig. 6). We conclude that SARS-CoV-2 antigen specific CD4^+^ T cells are induced during acute infection and are only slightly reduced after 6.1 months.

**Fig. 3.**
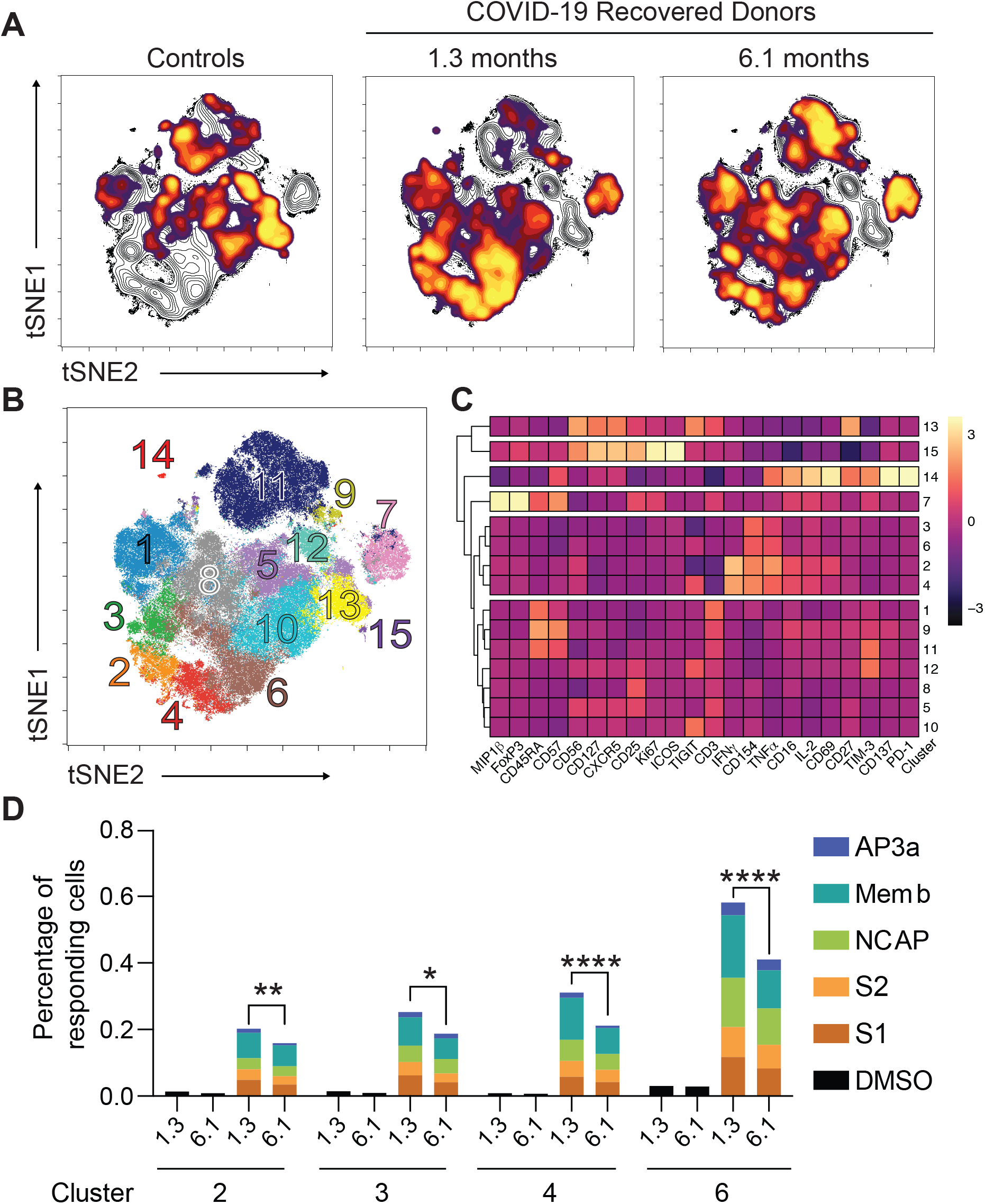
Antigen-specific CD4^+^ T cells dynamics in COVID-19 convalescent individuals. (**A**) viSNE representations of CD137^+^ CD154^+^ SARS-CoV-2-stimulated CD4^+^ T cells in unexposed individuals (controls) and COVID-19 convalescent individuals pooled. Density plots from each group concatenated is overlaid on the total contour viSNE plot. (**B**) viSNE representation of antigen-specific CD4^+^ T cell clusters, identified by FlowSOM clustering. (**C**) Column-scaled *z*-scores of median fluorescence intensity (MFI) as indicated by cluster and marker. (**D**) Percentage of antigen-specific CD4^+^ cells in the indicated FlowSOM clusters. Each bar represents the mean percentage for all COVID-19 convalescent individuals for the indicated SARS-CoV-2 peptide pools. *p<0.05, **p<0.01, ***p<0.001, ****p<0.0001.

To characterize SARS-CoV-2 antigen specific cytokine producing CD4^+^ T cells we analyzed the high dimensional flow cytometry data by traditional gating (Fig. 4 and Suppl. Fig. 2). IL-2, INFγ, and TNFα responses to peptide pools corresponding to Spike (S1 and S2), NCAP, Memb, AP3a, and CMV control were measured independently at both time points (Fig. 4A-C). In all cases, responses to the individual peptide pools were elevated above control at both time points, and all but NCAP and Memb responses remained stable between the time points (Fig. 4A-C). Among the individuals tested 97.5% and 95% responded to at least one of the antigens at 1.3 and 6.1 months respectively (Suppl. Fig. 7A-C). When all cytokines are considered together, we find a significant increase in antigen specific CD4^+^ T cell responses to all of the individual peptide pools at both time points compared to control (Fig. 4A). The increase in CD4^+^ T cell cytokine responses to the individual peptide pools and the overall combination of all SARS-CoV-2 antigens was not driven by any single cytokine but instead reflected increases in each of the three cytokines measured (Suppl. Fig. 7D). Moreover, when all antigen specific responses are considered in aggregate, the fraction of responding CD4^+^ T cells remains significantly elevated and is not different between the 2 time points (Fig. 4A, All). Antigen specific memory CD4^+^ memory T cells (CD45RA^+^CD27^+^) were similarly elevated at the two time points (Fig. 4B). Finally, there was no significant change in the CD4^+^ T cell response by the same individuals to the control CMV peptide pool (Fig. 4A-C).

**Fig. 4.**
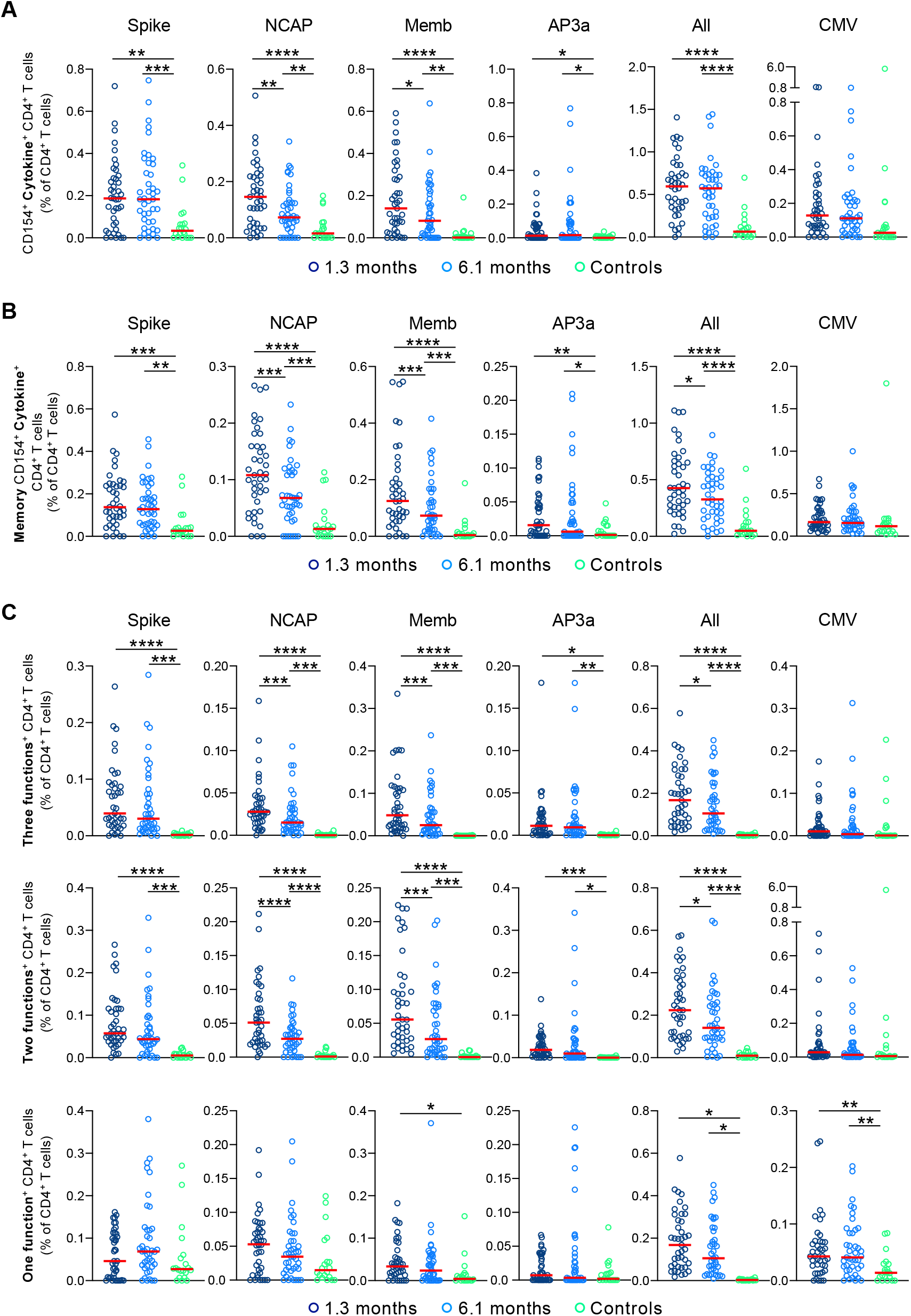
SARS-CoV-2-specific CD4^+^ T cells responses in convalescent COVID-19 individuals. Longitudinal analysis of COVID-19 specific CD4^+^ T cell responses in paired samples obtained 1.3 months and 6.1 months after infection. Cytokine production (IL-2, IFN-γ, and TNF-α) by CD4^+^ T cells analyzed by Intracellular Cytokine Staining (ICS). Spike (Aggregation of responses to Spike peptide pool S1 and S2), Nucleocapsid (NCAP) Membrane (Memb) and non-structural protein 3a (AP3a) peptide pools responses by controls (n=20), and convalescent COVID-19 individuals (n=41). **(A)** Combined frequency of SARS-CoV-2 specific CD4^+^ T cells that produce either IL-2, IFN-γ, and/or TNF-α. **(B)** Frequency of SARS-CoV-2 specific memory CD4^+^ T cells (CD45RA^-^ CD27^+^) that produce either IL-2, IFN-γ, and/or TNF-α. (**C**) Frequency of SARS-CoV-2-specific CD4^+^ T cells that produce either 3 cytokines, 2 cytokines or 1 cytokine. Each dot represents an individual at 1.3 months (dark blue) or 6.1 months (light blue) or control individuals (green). Significance determined by paired t test for comparisons between time points within individuals and unpaired t test for comparison between controls and COVID-19 individuals. *p<0.05, **p<0.01, ***p<0.001, ****p<0.0001.

Polyfunctional cytokine responses are associated with effective cellular immune responses (Seder et al., 2008, Lin et al., 2015, Betts et al., 2006). Polyfunctional CD4^+^ T cells responses to each of the individual peptide pools were significantly elevated at the early time point and remained so after 6.1 months (Fig. 4C). The magnitude of these responses was directly correlated with antibodies to the SARS-CoV-2 receptor binding domain (Suppl. Fig. 8). However, there was a decrease in polyfunctional CD4^+^ T cell responses to NCAP and Memb antigens at 6.1 months which is also reflected in a 22% decrease in the overall tri-functional response to the combined SARS-CoV-2 peptide libraries (Fig. 4C, All). In contrast, CD4^+^ T cells that produced only a single cytokine were only elevated in response to Memb and only at the early time point (Fig. 4C). We conclude that robust polyfunctional CD4^+^ T cell responses persist for 6.1 months after SARS-CoV-2 infection but decrease significantly when compared to an earlier time point.

### SARS-CoV-2 antigen specific CD8^+^ T cells

To examine antigen specific CD8^+^ T cell responses we measured production of Mip-1β, CD107a, IL-2, INF-γ and TNF-α in response to stimulation with SARS-CoV-2 peptide pools *in vitro*. In contrast to CD4^+^ T cells, CD8^+^ T cell responses were far more variable and generally less robust making tSNE analysis less reliable and therefore these responses were only analyzed by traditional gating (Fig. 5 and Suppl. Fig. 9). Although 95% of the donors tested responded to at least 1 of the peptide pools at both 1.3 and 6.1 months, the percentage of responding cells was low (Fig. 5A). When all peptide responses were pooled, we found significant Mip-1β, and INF-γ responses at both time points, while CD107a was only increased above control at 1.3 months. Polyfunctional responses that included at least 3 different cytokines were also elevated at both time points (Fig. 5B). We conclude that although we detect fewer antigen specific CD8^+^ than CD4^+^ T cells to SARS-CoV-2, they persist 6.1 months after infection.

**Fig. 5.**
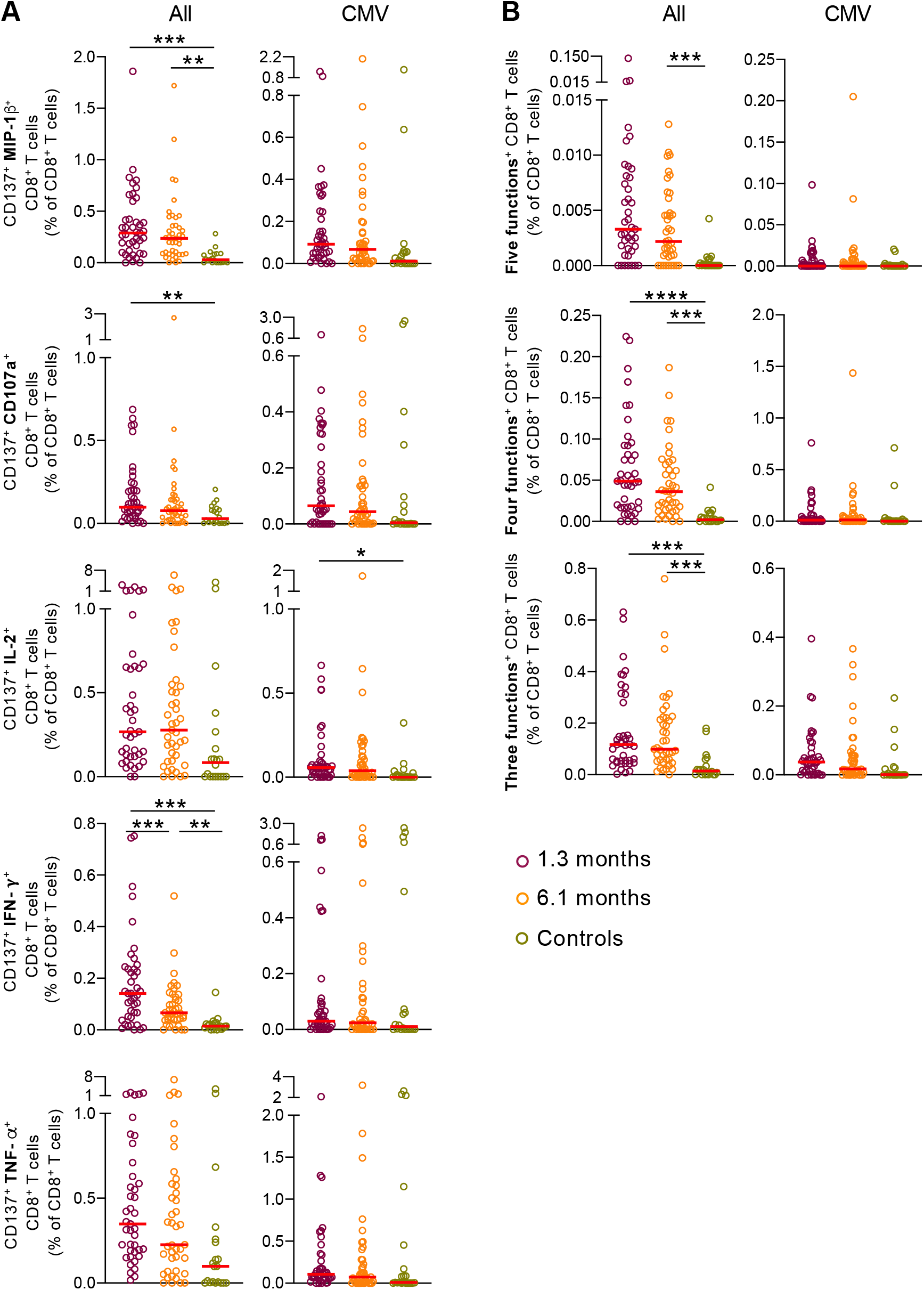
SARS-CoV-2-specific CD8^+^ T cells responses in convalescent COVID-19 individuals. Longitudinal analysis of COVID-19 specific CD8^+^ T cell responses in paired samples obtained from the same individual at 1.3 months and 6.1 months after infection. Cytokine production (MIP-1β, CD107a, IL-2, IFN-γ, and TNF-α) by CD8^+^ T cells analyzed by Intracellular Cytokine Staining (ICS). Spike (Aggregation of responses to spike peptide pool S1 and S2), Nucleocapsid (NCAP) Membrane (Memb) and non-structural protein 3a (AP3a) peptide pools responses by controls (n=20), and convalescent COVID-19 individuals (n=41). **(A)** Frequency of SARS-CoV-2 specific CD8^+^ T cells that produce MIP-1β, CD107a, IL-2, IFN-γ, or TNF-α. **(B)** Frequency of SARS-CoV-2 specific CD8^+^ T cells that produce either 5 cytokines, 4 cytokines or 3 cytokines. Each dot represents an individual at 1.3 months (dark red) or 6.1 months (orange) or control individuals (green). Significance determined by paired t test for comparisons between time points within individuals and unpaired T test for comparison between controls and COVID-19 individuals. *p<0.05, **p<0.01, ***p<0.001, ****p<0.0001.

## Discussion

Most effective vaccines and anamnestic responses to pathogens are mediated by neutralizing antibodies (Plotkin, 2020, Inoue et al., 2018, Weisel and Shlomchik, 2017). Consistent with this notion, neutralizing antibodies are protective against SARS-CoV-2 infection in animal models (Deng et al., 2020) and appear to correlate with protection in vaccinated humans (Gaebler and Nussenzweig, 2020). However, antibody responses are dependent on specialized helper T cells that control the activation and selection of antibody producing plasma and memory B cells (Victora and Nussenzweig, 2012, Crotty, 2015).

CD4^+^ and CD8^+^ T cells can also contribute directly to protection against SARS-CoV (Le Bert et al., 2020, Tang et al., 2011, Yang et al., 2006, Channappanavar et al., 2014) and other viral pathogens in animal models (Schmitz et al., 1999, Shoukry et al., 2003, Snyder, 2011). In addition, control of human viral pathogens such as HIV-1 is associated with CD8^+^ T cells in rare elite controllers (Collins et al., 2020). To gain further understanding into whether SARS-CoV-2 infection is associated with enduring responses we examined cellular immunity to SARS-CoV-2 in paired samples collected 1.3 and 6.1 months after infection. Similar to SARS-CoV and MERS infections, SARS-CoV-2 antigen specific memory CD4^+^ and CD8^+^ T cells persist and would be expected to play role in protection against re-exposure.

In addition to persistent antigen specific memory responses, chronic viral infections such as HIV are associated with lasting immune perturbations (Breton et al., 2013), but little is known about acute viral infections. SARS-CoV-2 is an acute infection that typically resolves after 2-3 weeks and in rare instances leads to severe lung disease and mortality (Richardson et al., 2020, O’Driscoll et al., 2020). The cohort we examined is biased toward milder forms of the disease, nevertheless, CD4^+^ and CD8^+^ T cell subset distribution, cell division and expression of activation/exhaustion markers remain altered 6 months after SARS-CoV-2 infection. These abnormalities were not directly associated with persistent symptoms and their impact on the overall immune health of the individual remains to be determined.

## Materials and Methods

### Study participants

Study participants (n = 61) were residents of the Greater New York City tri-state region, 20 of whom were SARS-CoV-2 unexposed (pre-pandemic, median age: 52.5 years old, 45% female), and 41 were SARS-CoV-2-infected (median age: 45 years old; 36.6% female). Previously enrolled study participants (Robbiani et al., 2020) were asked to return for a 6-month follow-up visit at the Rockefeller University Hospital in New York from August 31 through October 16, 2020. All of the individuals tested had RT-PCR confirmed SARS-CoV-2 infection or were close contacts that seroconverted. A summary of the participants’ clinical characteristics is presented in Table 1 and has been extensively described elsewhere (Robbiani et al., 2020, Gaebler et al., 2020).

### Cell preparation

Blood samples were collected an average of 1.3 and 6.1 months post infection (Table 1). Peripheral blood mononuclear cells (PBMCs) were isolated from heparinized blood by density gradient centrifugation (Ficoll-Paque™) and cryopreserved in 90% heat-activated fetal bovine serum (FBS) plus 10% DMSO in liquid nitrogen. Thawed PBMCs were washed and resuspended at 2×10^6^ cells/ml with RPMI 1640 supplemented with 10% heat inactivated human serum (GemCell) and 10 U/ml Benzonase. For cell stimulation experiments cells were rested at 37°C and 5% CO_2_ for 8 hours prior to stimulation with peptides for use in intracellular cytokine staining assays.

### Synthetic COVID-19 peptides

Pool of 315 peptides (15mers with 11 aa overlap) (Spike glycoprotein, delivered in two subpools of 158 (S1 pool) & 157 (S2 pool) peptides), 102 peptides (Nucleoprotein), 53 peptides (Membrane protein) and 66 peptides (Protein 3a, AP3a) of SARS-CoV-2 were purchased from JPT (Berlin, Germany). Pool of 138 peptides (15mers with 11 aa overlap) derived from 65 kDa phosphoprotein (pp65) of Human cytomegalovirus (CMV) (JPT) was used as control. Staphylococcus Enterotoxin B (SEB) (Sigma-Aldrich, St-Louis, MO, USA) was used at a final concentration of 1μg/ml as a positive stimulation control. Peptides were reconstituted in high grade DMSO (Sigma-Aldrich, St-Louis, MO, USA) at a concentration of 0.1 mg/ml and used at a final concentration of 0.25 μg/ml in a maximum of 0.2% DMSO. PBMCs with peptide diluent (0.2% DMSO) served as the negative control.

### Cell Stimulation and intracellular staining

Cells were stimulated for 12h with SARS-CoV-2 peptide pools (0.25 μg/ml) in the presence of αCD28/αCD49d co-stimulatory antibodies (BD FastImmune™, BD Biosciences, San Diego, CA, USA), 5 μg/ml brefeldin A (Sigma-Aldrich, St-Louis, MO, USA) and 5 μg/ml Monensin. (BD GolgiStop™ BD Biosciences, San Diego, CA, USA), and anti-CD107a-PE-Cy5 (Clone H4A3) (the staining for CD107a is carried out during cell activation in this assay). A negative control containing PBMCs and co-stimulatory antibodies from the same subject, with DMSO, was also included for each assay. Following stimulation, cells were washed with PBS and were first stained with CCR7 (Clone 3D12) at 37°C for 20 min in PBS containing 2% FBS. Cells were then surface stained for 30 min in the dark at 4°C with viability reagent (BD Horizon™ Fixable Viability Stain, BD Biosciences, San Diego, CA, USA) and a 29-color cocktail of monoclonal antibodies (mAbs) containing surface antibodies against CD19 (Clone SJ25C1), CD20 (Clone 2H7), CD66b (Clone G10F5), CD14 (Clone M5E2), CD16 (Clone 3G8), CD56 (Clone B159), CD4 (Clone OKT4), CD8 (Clone RPA-T8), CD45RA (Clone HI100), CD27 (Clone M-T271), CD57 (Clone NK-1), CD25 (Clone 2A3), CXCR5 (Clone RF8B2), CD127 (Clone HIL-7R-M21), PD-1 (Clone EH12.1), TIGIT (Clone 741182), Tim-3 (Clone 7D3) and CD69 (Clone FN50). The cells were then washed with PBS containing 2% FBS and permeabilized according to the manufacturer’s instructions using a Foxp3/Transcription Factor Staining Buffer Set (eBioscience™) and stained with intracellular antibodies against CD3 (Clone SK7), MIP-1b (Clone D21-1351), IL-2 (Clone MQ1-17H12), IFN-g (Clone B27), TNF-a (Clone MAb11), FoxP3 (Clone 236A/E7), Ki67 (Clone ki-67), CD154 (Clone 24-3) and CD137 (Clone 4B4-1). After labelling, cells were washed and fixed in PBS containing 2% paraformaldehyde and stored at 4°C prior to flow cytometry acquisition within 24 hours.

### Phenotypic characterization and surface staining

Cells from all donors were initially stained with CCR7 (Clone 3D12) at 37°C for 20 min in PBS containing 2% FBS. Cells were then stained with a 29-color staining cocktail containing viability reagent and surface antibodies against CD19 (Clone SJ25C1), CD20 (Clone 2H7), CD66b (Clone G10F5), CD14 (Clone M5E2), CD16 (Clone 3G8), CD56 (Clone B159), CD3 (Clone SK7), CD4 (Clone OKT4), CD8 (Clone RPA-T8), CD45RA (Clone HI100), CD27 (Clone M-T271), CD95 (Clone DX2), NKG2A (Clone 131411), NKG2C (Clone 134591), PD-1 (Clone EH12.1), TIGIT (Clone 741182), TIM-3 (Clone 7D3), CD152 (Clone BNI3), CD272 (Clone J168-540), CD134 (Clone ACT35), CD274 (Clone MIH1), CD273 (Clone MIH18), CD96 (Clone 6F9), CD357 (Clone V27-580), CD160 (Clone BY55), CD137 (Clone 4B4-1), CD223 (Clone T47-530), CD278 (Clone C398.4A), CD28 (Clone CD28.2) and CD244 (Clone eBioC1.7). Titrated antibodies were added to 2 million cells in 50 μl of PBS containing 2% FBS for 30 minutes at 4°C. Washed cells were then fixed in 2% formaldehyde and stored at 4°C until analysis.

### Flow cytometry analysis

All events -approximately 1,200,000 to 1,800,000 events per sample-were collected on a BD FACSymphony™A5 Cell Analyzer (BD Biosciences, San Diego, CA, USA). The lymphocytes were gated for further analysis, as described in Figure S2, using Flowjo™ Software Version 9.9.6 (BD, USA). For cytokine expression, we subtracted the background in the negative control. To ensure equivalent fluorescence intensities (median fluorescence intensity or MFI) from one experiment to another, we used Rainbow beads (Spherotech, Inc, Lake Forest, IL, USA). For every run, rainbow beads were acquired first and the voltages were adjusted if necessary to accommodate both daily variations and larger changes in performance such as would be seen after cytometer maintenance and alignment.

### High-dimensional data analysis of flow cytometry data

viSNE and FlowSOM analyses were performed on Cytobank (https://cytobank.org). viSNE analysis was performed using equal sampling of 5000 cells (for total T cell phenotyping analysis, Fig. 1), or proportional sampling (for antigen-specific CD4^+^ T cell analysis, Fig. 3) from each FCS file, with 7500 iterations, a perplexity of 30, and a theta of 0.5. The following markers were used to generate viSNE maps: CD273, CD96, NKG2C, CD152, TIGIT, CD272, CD134, CD45RA, CD137, CD366, CD95, CD279, CD16, CD274, CD27, CD56, CD357, CCR7, CD223, CD278, CD28, CD244, NKG2a, CD4, and CD8 (for total T cells, Fig. 1), and CXCR5, CD127, MIP1b, CD137, IL-2, TIGIT, CD45RA, CD57, CD25, TIM-3, CD69, PD-1, CD16, IFNg, CD27, CD56, TNFa, Ki67, ICOS, CD107a, FoxP3, CD154 (for antigen-specific CD4^+^ T cells, Fig. 3 and Suppl. Figs. 5-6). Resulting viSNE maps were fed into the FlowSOM clustering algorithm (Van Gassen et al., 2015). The self-organizing map (SOM) was generated using hierarchical consensus clustering on the tSNE axes.

### Statistical analysis

Statistical analyses were performed using Prism 7.0 (GraphPad). Significances between matched groups were calculated using paired t-test whereas differences between unmatched groups were compared using unpaired t-test, or one-way ANOVA. Correlations were performed using the Spearman rank correlation test.

## ACKNOWLEDGMENTS

We thank all study participants who devoted time to our research; Drs. Barry Coller and Sarah Schlesinger, the Rockefeller University Hospital Clinical Research Support Office and nursing staff. All members of the M.C.N. laboratory for helpful discussions and Maša Jankovic for laboratory support.

## Author contributions

G.B. and M.C.N. conceived and designed the study. G.B., P.M. and T.H. carried out and analyzed the experiments. D.S.B., M.C. and C.G. designed clinical protocols. M.C., M.T. and A.H. recruited participants and executed clinical protocols. T.Y.O. advised on bioinformatic analysis. G.B., P.M, T.H. and M.C.N. wrote the manuscript with input from all co-authors.

## Competing interests

None.

**Suppl. Fig. 1. Persistent longitudinal changes in the phenotypic landscape of T cells in individuals recovered from COVID-19, related to Figure 1**. (**A**) viSNE projections of the indicated protein expression. (**B**) Frequency of T cells from each group in FlowSOM clusters indicated.

**Suppl. Fig. 2**. Gating strategy to identify CD4^+^ and CD8^+^ T cells and their subsets.

**Suppl. Fig 3. Phenotypic characteristics of central memory and cycling T cells**. (**A**) PD-1, TIGIT and TIM-3 expression of CD4^+^ central memory T cells. (**B**) PD-1, TIGIT and TIM-3 expression of CD8^+^ central memory T cells. (**C**) PD-1, TIGIT, TIM-3 and CD25 expression of CD4^+^ cycling T cells (Ki67^+^). (**D**) PD-1, TIGIT, TIM-3 and CD25 expression of CD8^+^ cycling T cells (Ki67^+^).

**Suppl. Fig. 4. Frequency of circulating T Follicular Helper cells (T**_**FH**_**) and T regulatory cells (Treg)**. (**A**) Circulating T Follicular Helper cells (cT_FH_) and (**B**) T regulatory cells (Treg) relative numbers. Each dot represents a COVID-19 convalescent individual at 1.3 months (dark blue) or 6.1 months (light blue) or control individuals (green). Significance determined by paired t test for comparisons between time points within individuals and unpaired T test for comparison between unexposed and COVID-19 individuals. *p<0.05, **p<0.01, ***p<0.001, ****p<0.0001.

**Suppl. Fig. 5. Antigen-specific CD4**^**+**^ **T cells dynamics responding to individual SARS-CoV-2 peptide pools in COVID-19 convalescent individuals, related to Figure 3**. (**A**) Mean fluorescence intensity (MFI) for indicated markers, column-normalized z-score. (**B**) viSNE representations of CD137^+^ CD154^+^ SARS-CoV-2-stimulated CD4^+^ T cells in unexposed individuals (controls) and COVID-19 convalescent individuals pooled. Density plots from each group concatenated is overlaid on the total contour viSNE plot.

**Suppl. Fig. 6. Antigen-specific CD4**^**+**^ **T cells dynamics to SARS-CoV-2 and CMV in COVID-19 convalescent individuals**. (**A**) viSNE representations of CD137^+^ CD154^+^ CD4^+^ T cells stimulated with SARS-CoV-2 or CMV in unexposed individuals (controls) and COVID-19 convalescent individuals pooled. Density plots from each group concatenated is overlaid on the total contour viSNE plot. (**B**) viSNE representation of each indicated marker expression. (**C**) viSNE representation of antigen-specific CD4^+^ T cell clusters, identified by FlowSOM clustering. (**D**) Mean fluorescence intensity (MFI) for indicated markers, column-normalized z-score. (**E**) Percentage of antigen-specific CD4^+^ cells in the indicated FlowSOM clusters. Each bar represents the mean percentage for all COVID-19 convalescent individuals for the indicated SARS-CoV-2 peptide pools or for CMV. *p<0.05, **p<0.01, ***p<0.001, ****p<0.0001.

**Suppl. Fig. 7. SARS-CoV-2-specific CD4**^**+**^ **T cells responses in convalescent COVID-19 individuals, related to Figure 3. (A)** Gating strategy for identification of SARS-CoV-2-specific CD4^+^ T cells. **(B)** Percentage of COVID-19 individuals that respond to Spike (aggregation of responses to spike peptide pool S1 and S2), Nucleocapsid (NCAP) Membrane (Memb) and non-structural protein 3a (AP3a) peptide pools at 1.3 months or 6.1 months. **(C)** Pie chart shows the frequency of recovered COVID-19 individuals that respond to either 1, 2, 3, 4 or 5 peptide pools. **(D)** Frequency of SARS-CoV-2-specific CD4^+^ T cells that produce either IL-2, IFN-γ or TNF-α. Each dot represents an individual with COVID-19 at 1.3 months (dark blue or 6.1 months (light blue) or control individuals (green). Significance determined by paired t test for comparisons between time points within individuals and unpaired T test for comparison between unexposed and COVID-19 individuals. *p<0.05, **p<0.01, ***p<0.001, ****p<0.0001.

**Suppl. Fig. 8. Correlations of polyfunctional CD4**^**+**^ **T cells with antibody titers**. Normalized AUC for IgG anti-RBD plotted against the relative frequency of CD4^+^ T cells producing 3 cytokines (three functions). The r and p values were determined by two-tailed Spearman’s correlations.

**Suppl. Fig. 9. SARS-CoV-2-specific CD8**^**+**^ **T cells responses in convalescent COVID-19 individuals, related to Figure 4. (A)** Gating strategy for identification of SARS-CoV-2-specific CD8^+^ T cells. **(B)** Percentage of COVID-19 individuals that respond to Spike (aggregation of responses to spike peptide pool S1 and S2), Nucleocapsid (NCAP) Membrane (Memb) and non-structural protein 3 (AP3) peptide pools at 1.3 months or 6.1 months. **(C)** Pie chart shows the frequency of mild COVID-19 individuals that have CD8^+^ responses to either 1, 2, 3, 4 or 5 peptide pools. **(D)** Frequency of SARS-CoV-2 specific CD4^+^ T cells that produce either IL-2, IFN-γ or TNF-α. **(E)** Frequency of SARS-CoV-2-specific CD8^+^ T cells that produce either 5 cytokines, 4 cytokines or 3 cytokines. Each dot represents an individual with COVID-19 at 1.3 months (dark red) or 6.1 months (orange) or unexposed individuals (green). Significance determined by paired t test for comparisons between time points within individuals and unpaired t test for comparison between unexposed and COVID-19 individuals. *p<0.05, **p<0.01, ***p<0.001, ****p<0.0001.

**Supplementary Table 1**: Individual participant characteristics.

