## Supplementary Figures for "Persistent Cellular Immunity to SARS-CoV-2 Infection"

Figure S1.

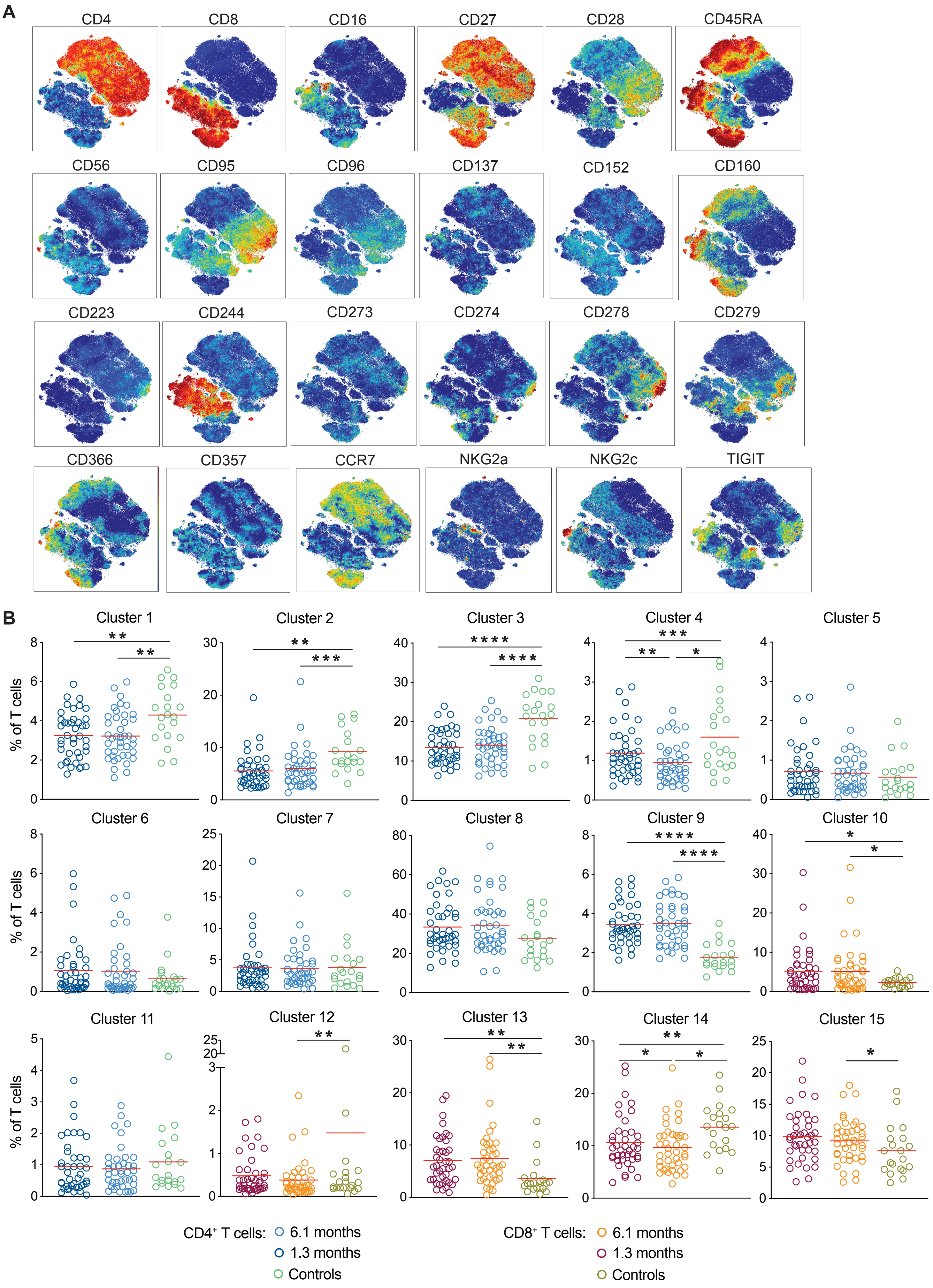
A
CD4
CD8
CD16
CD27
CD28
CD45RA
CD56
CD95
CD96
CD137
CD152
CD160
CD223
CD244
CD273
CD274
CD278
CD279
CD366
CD357
CCR7
NKG2a
NKG2c
TIGIT
B
Cluster 1
Cluster 2
Cluster 3
Cluster 4
Cluster 5
Cluster 6
Cluster 7
Cluster 8
Cluster 9
Cluster 10
Cluster 11
Cluster 12
Cluster 13
Cluster 14
Cluster 15
% of T cells
CD4⁺ T cells: 6.1 months
1.3 months
Controls
CD8⁺ T cells: 6.1 months
1.3 months
Controls

Figure S2.

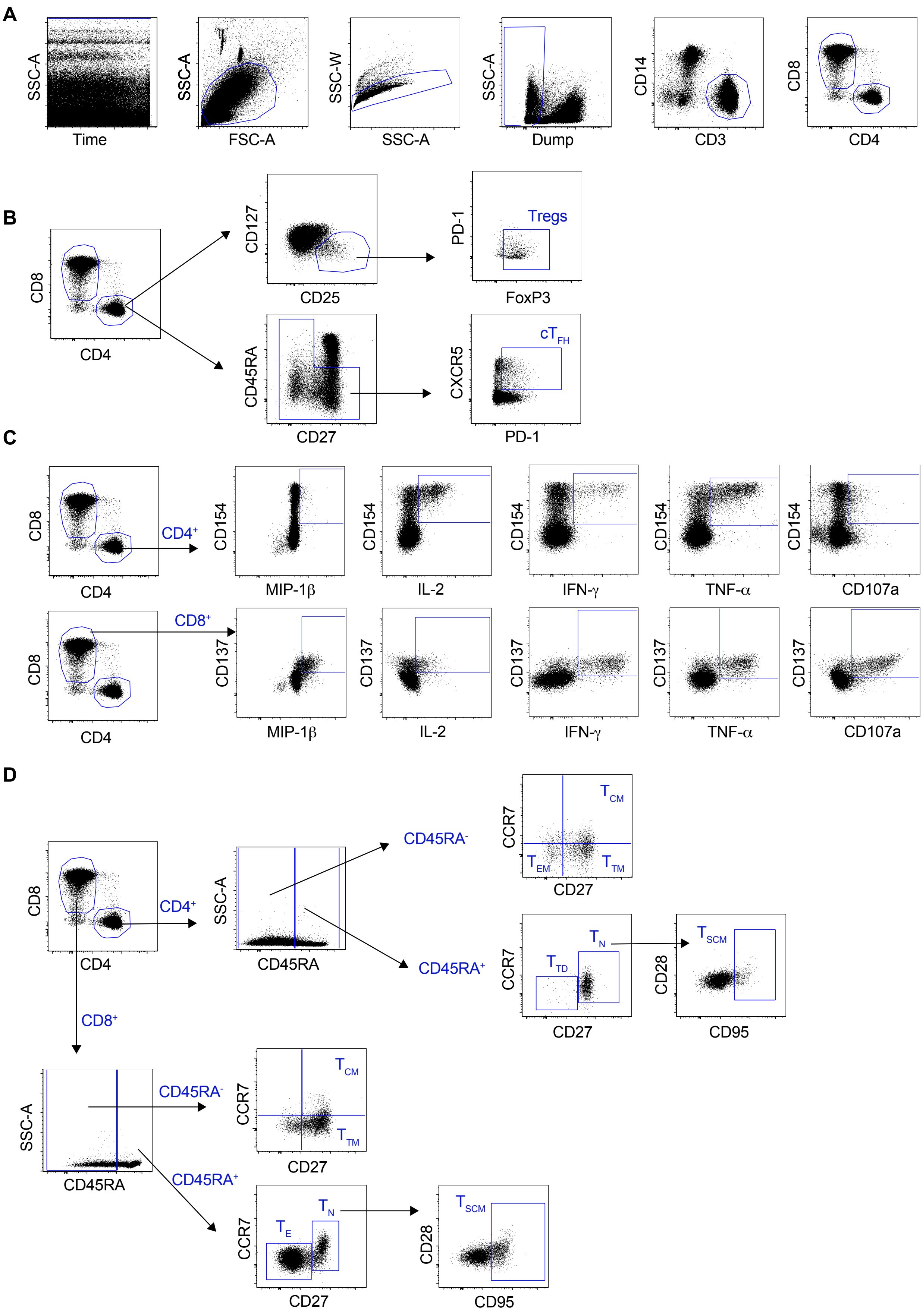
A
SSC-A
Time
SSC-A
FSC-A
SSC-W
SSC-A
SSC-A
Dump
CD14
CD3
CD8
CD4
B
CD8
CD4
CD127
CD25
PD-1
FoxP3
Tregs
CD45RA
CD27
CXCR5
PD-1
cT FH
C
CD8
CD4
CD4+
CD154
MIP-1β
CD154
IL-2
CD154
IFN-γ
CD154
TNF-α
CD154
CD107a
CD8
CD4
CD8+
CD137
MIP-1β
CD137
IL-2
CD137
IFN-γ
CD137
TNF-α
CD137
CD107a
D
CD8
CD4
CD4+
SSC-A
CD45RA
CD45RA-
CD45RA+
CCR7
CD27
T CM
T EM
T TM
CCR7
CD27
T N
T TD
CD28
CD95
T SCM
CD8+
SSC-A
CD45RA
CD45RA-
CD45RA+
CCR7
CD27
T CM
T TM
CCR7
CD27
T N
T E
CD28
CD95
T SCM

Figure S3.

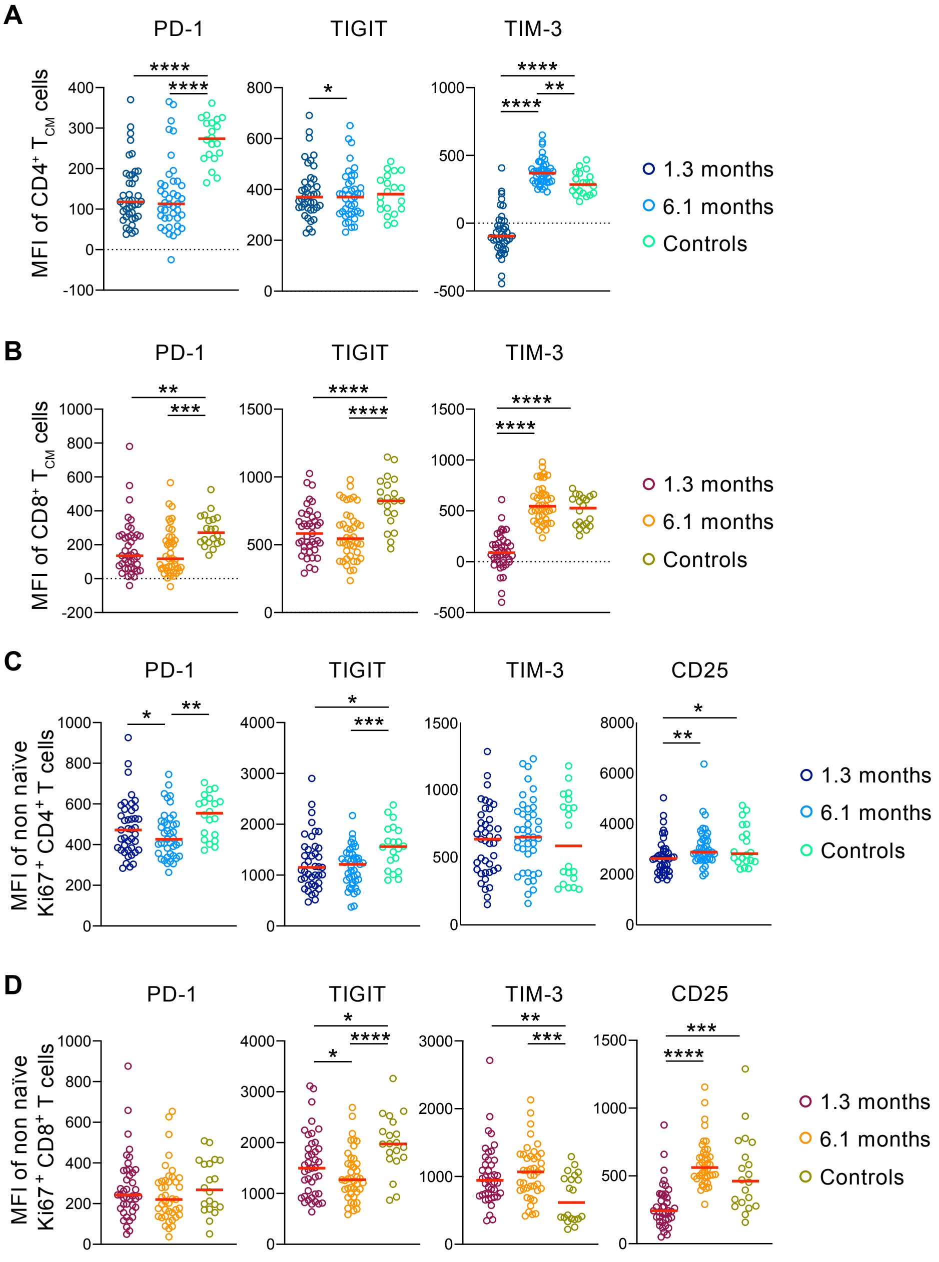
A
PD-1
TIGIT
TIM-3
MFI of CD4+ TCM cells
1.3 months
6.1 months
Controls
B
PD-1
TIGIT
TIM-3
MFI of CD8+ TCM cells
1.3 months
6.1 months
Controls
C
PD-1
TIGIT
TIM-3
CD25
MFI of non naïve Ki67+ CD4+ T cells
1.3 months
6.1 months
Controls
D
PD-1
TIGIT
TIM-3
CD25
MFI of non naïve Ki67+ CD8+ T cells
1.3 months
6.1 months
Controls

Figure S4.

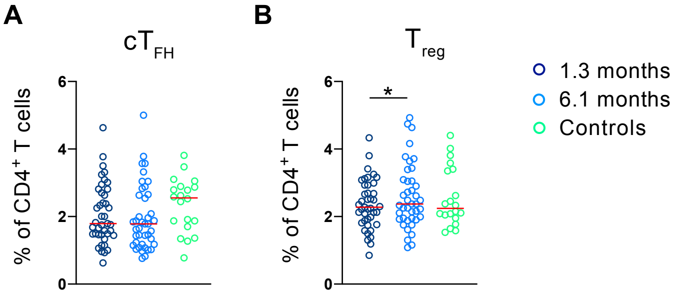
A
cTFH
% of CD4+ T cells
6
4
2
0
B
Treg
*
% of CD4+ T cells
6
4
2
0
1.3 months
6.1 months
Controls

Figure S5.

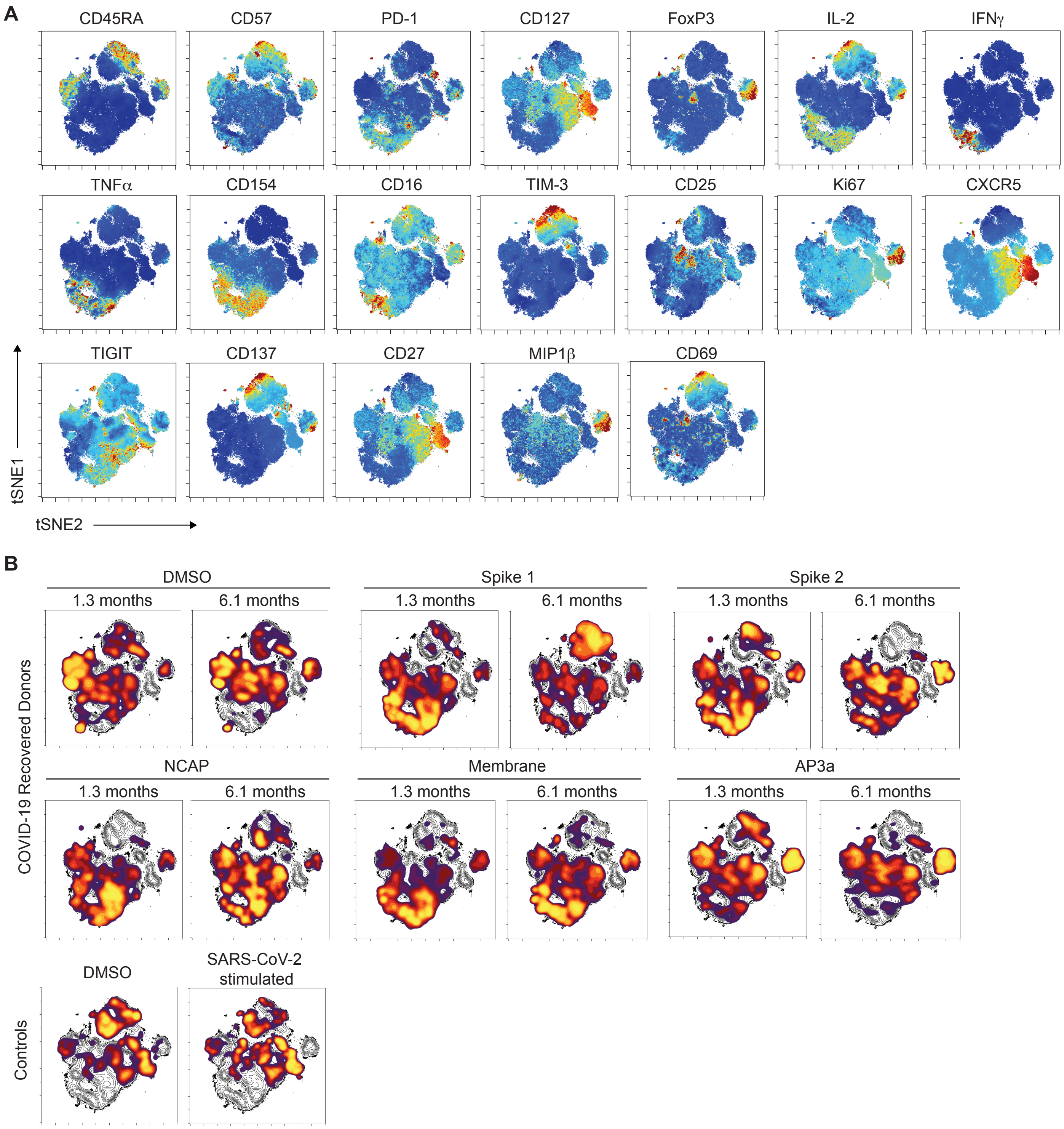

A
CD45RA
CD57
PD-1
CD127
FoxP3
IL-2
IFNγ
TNFα
CD154
CD16
TIM-3
CD25
Ki67
CXCR5
TIGIT
CD137
CD27
MIP1β
CD69
tSNE1
tSNE2
B
DMSO
Spike 1
Spike 2
1.3 months
6.1 months
NCAP
Membrane
AP3a
COVID-19 Recovered Donors
Controls
DMSO
SARS-CoV-2 stimulated

Fig. S6

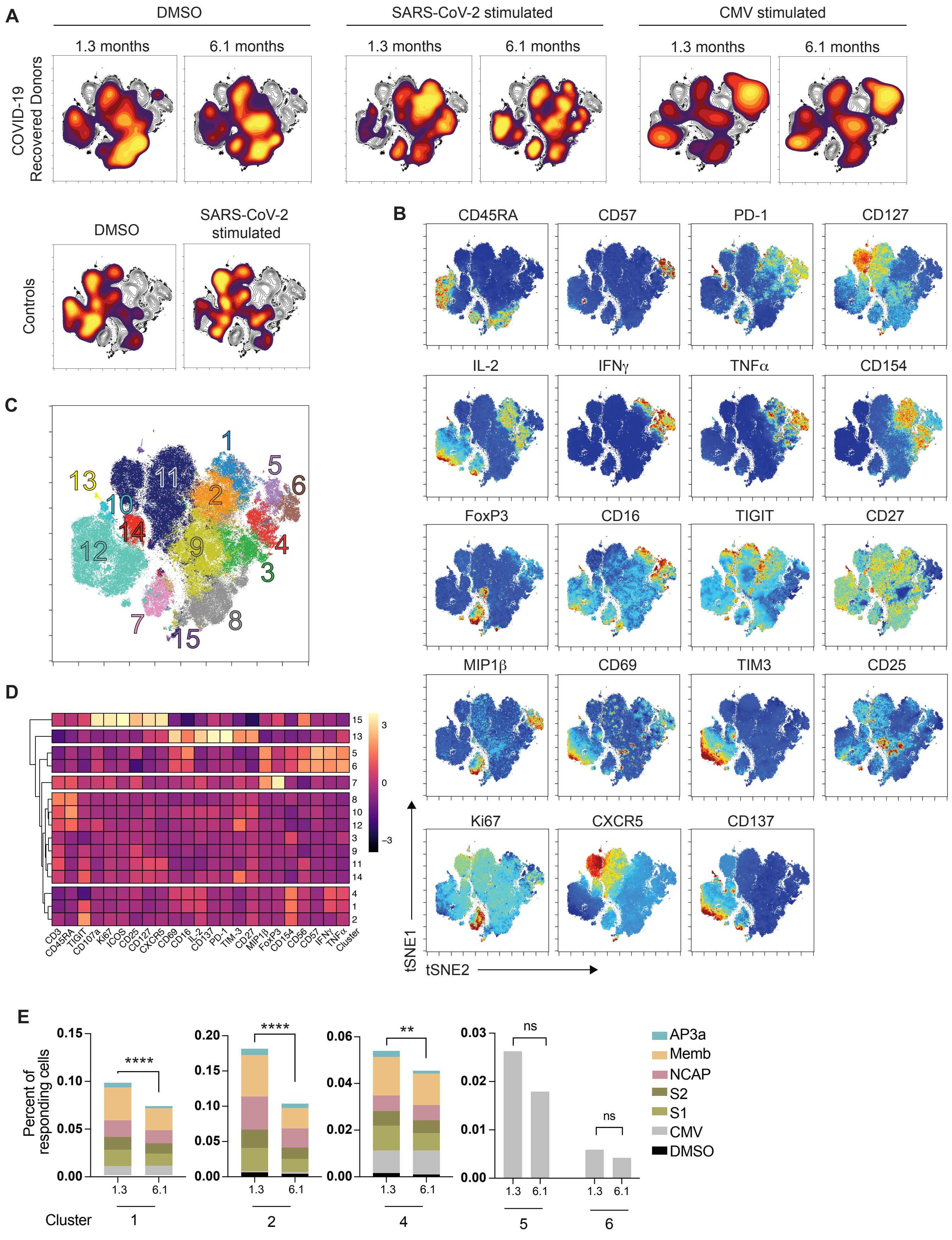
A
DMSO
SARS-CoV-2 stimulated
CMV stimulated
1.3 months
6.1 months
COVID-19 Recovered Donors
Controls
DMSO
SARS-CoV-2 stimulated
B
CD45RA
CD57
PD-1
CD127
IL-2
IFNγ
TNFα
CD154
FoxP3
CD16
TIGIT
CD27
MIP1β
CD69
TIM3
CD25
Ki67
CXCR5
CD137
tSNE1
tSNE2
C
1 2 3 4 5 6 7 8 9 10 11 12 13 14 15
D
CD3 CD45RA TIGIT CD107a Ki67 ICOS CD25 CD127 CXCR5 CD69 CD16 IL-2 CD137 PD-1 TIM-3 CD27 MIP1β FoxP3 CD154 CD56 CD57 IFNγ TNFα Cluster
15 13 5 6 7 8 10 12 3 9 11 14 4 1 2
3 0 −3
E
Percent of responding cells
**** **** ** ns ns
1.3 6.1
Cluster 1 2 4 5 6
AP3a
Memb
NCAP
S2
S1
CMV
DMSO

Figure S7.

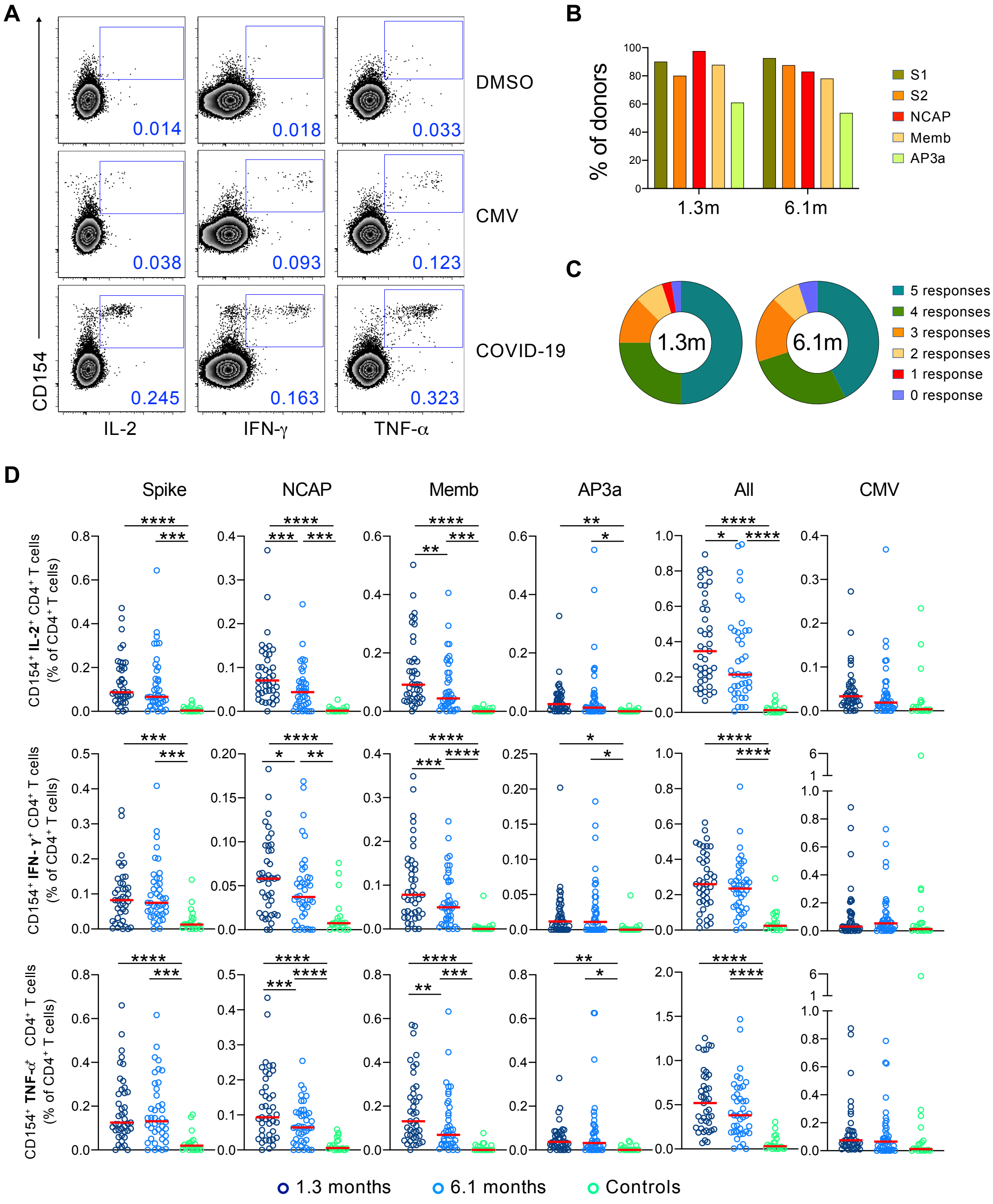
A
DMSO
CMV
COVID-19
0.014
0.018
0.033
0.038
0.093
0.123
0.245
0.163
0.323
CD154
IL-2
IFN-γ
TNF-α
B
% of donors
1.3m
6.1m
S1
S2
NCAP
Memb
AP3a
C
1.3m
6.1m
5 responses
4 responses
3 responses
2 responses
1 response
0 response
D
Spike
NCAP
Memb
AP3a
All
CMV
CD154+ IL-2+ CD4+ T cells (% of CD4+ T cells)
CD154+ IFN-γ+ CD4+ T cells (% of CD4+ T cells)
CD154+ TNF-α+ CD4+ T cells (% of CD4+ T cells)
1.3 months
6.1 months
Controls

Figure S8.

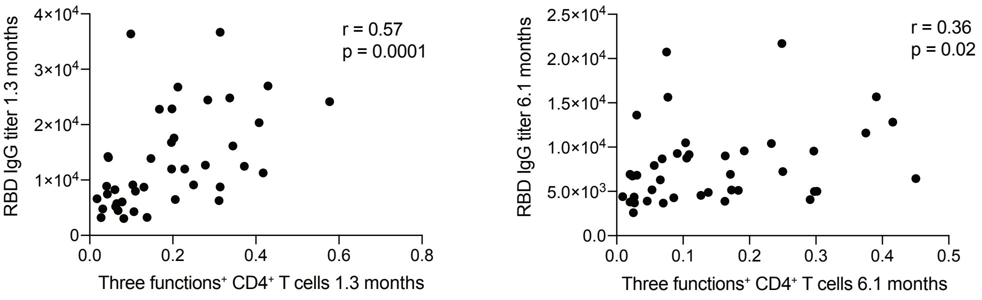

r = 0.57
p = 0.0001
RBD IgG titer 1.3 months
$4\times10^4$
$3\times10^4$
$2\times10^4$
$1\times10^4$
0
0.0
0.2
0.4
0.6
0.8
Three functions$^+$ CD4$^+$ T cells 1.3 months
r = 0.36
p = 0.02
RBD IgG titer 6.1 months
$2.5\times10^4$
$2.0\times10^4$
$1.5\times10^4$
$1.0\times10^4$
$5.0\times10^3$
0.0
0.0
0.1
0.2
0.3
0.4
0.5
Three functions$^+$ CD4$^+$ T cells 6.1 months

Figure S9.

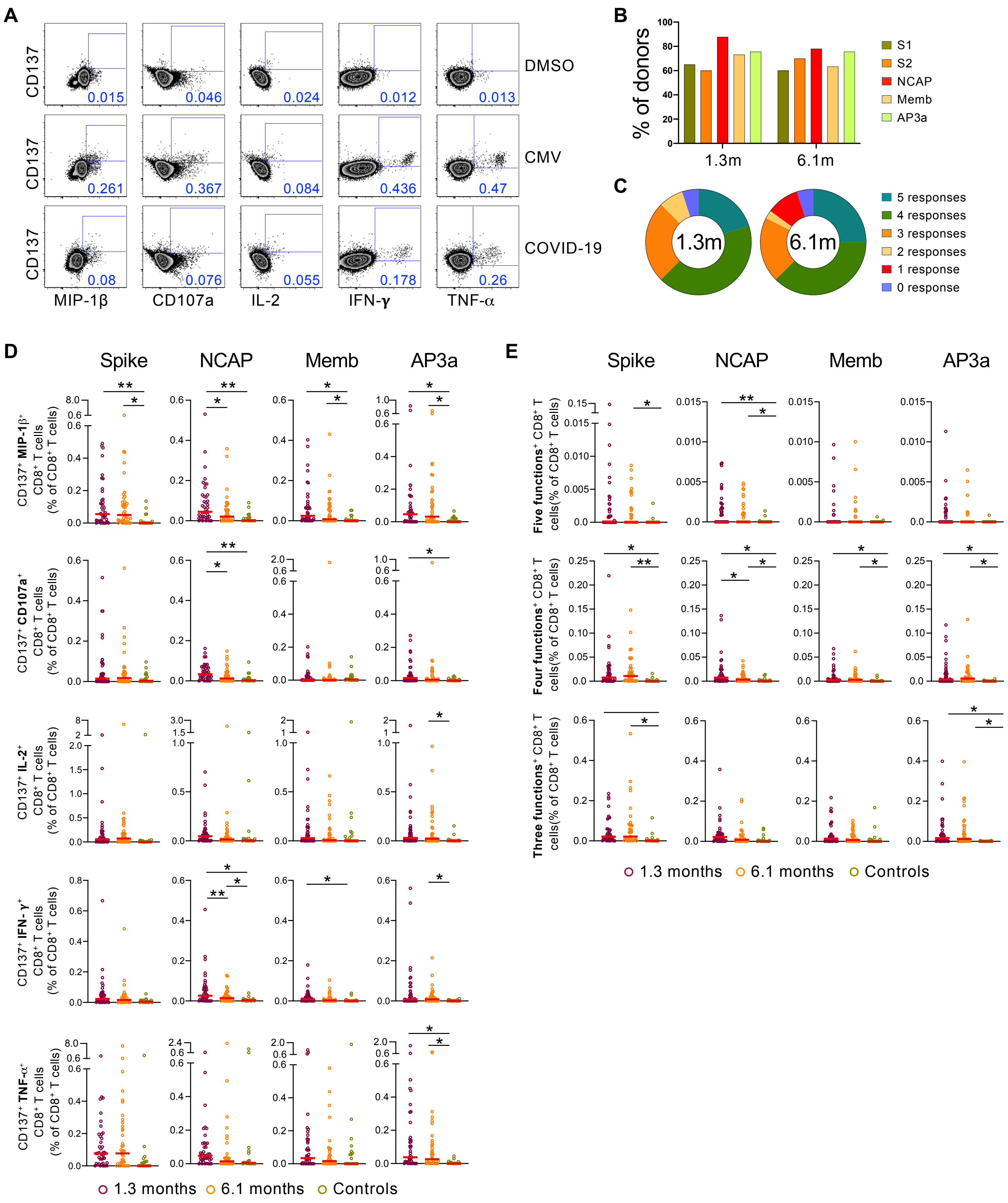
A
B
C
D
E
DMSO
CMV
COVID-19
MIP-1β
CD107a
IL-2
IFN-γ
TNF-α
% of donors
S1
S2
NCAP
Memb
AP3a
1.3m
6.1m
5 responses
4 responses
3 responses
2 responses
1 response
0 response
Spike
NCAP
Memb
AP3a
1.3 months
6.1 months
Controls

**Table 1. Individual participant characteristics**

| ID | Age (years) | Sex | Race | Ethnicity | Temporal dynamics (days): Sx duration in acute disease | Temporal dynamics (days): Sx onset to initial visit (T1) | Temporal dynamics (days): Sx onset to follow-up visit (T2) | Temporal dynamics (days): Time between visits | # of solicited comorbidities § | Acute disease severity by WHO (0-8) ¶ | Post-acute Sx persistence † | Serological assays: RBD (AUC) IgG (T1) | RBD (AUC) IgG (T2) | RBD (AUC) IgM (T1) | RBD (AUC) IgM (T2) | RBD (AUC) IgA (T1) | RBD (AUC) IgA (T2) | RBD (Pylon, IV) IgG (T1) | RBD (Pylon, IV) IgG (T2) | RBD (Pylon, IV) IgM (T1) | RBD (Pylon, IV) IgM (T2) | N (COI) total Ig (T1) | N (COI) total Ig (T2) | Neutralization (NT50) (T1) | Neutralization (NT50) (T2) |
|---|---|---|---|---|---|---|---|---|---|---|---|---|---|---|---|---|---|---|---|---|---|---|---|---|---|
| **7** | 40 | M | White | Non-Hispanic | 11 | 30 | 181 | 151 | 0 | 2 | Y | 11981 | 9545 | 6524 | 1516 | 1479 | 1344 | 34 | 18 | 3.6 | 0.31 | 56 | 171 | 2730 | 192 |
| **21** | 54 | M | White | Hispanic | 11 | 27 | 200 | 173 | 1 | 2 | Y | 36389 | 20744 | 14506 | 1242 | 2855 | 1914 | 127 | 59 | 4.4 | 0.23 | 82 | 159 | 5053 | 561 |
| **24** | 34 | M | White | Non-Hispanic | 15 | 30 | 175 | 145 | 1 | 1 | N | 5736 | 3803 | 2715 | 1150 | 927 | 1101 | 2.9 | 0.93 | 0.34 | 0.2 | 6 | 17 | 281 | 10 |
| **31** | 51 | M | White | Non-Hispanic | 9 | 33 | 183 | 150 | 0 | 2 | Y | 3212 | 3705 | 1272 | 903 | 906 | 913 | 1.3 | 1.5 | 0.22 | 0.15 | 35 | 41 | 192 | 18 |
| **46** | 39 | M | White | Non-Hispanic | 8 | 30 | 174 | 144 | 0 | 2 | Y | 4799 | 4416 | 2247 | 1315 | 1055 | 1153 | 3 | 2.2 | 0.24 | 0.17 | 51 | 141 | 59 | 21 |
| **47** | 43 | F | White | Non-Hispanic | 11 | 33 | 177 | 144 | 0 | 2 | Y | 17581 | 9284 | 9749 | 1914 | 1586 | 851 | 43 | 16 | 1.2 | 0.21 | 103 | 101 | 10433 | 349 |
| **48** | 37 | F | White | Non-Hispanic | 7 | 21 | 174 | 153 | 0 | 2 | N | 3265 | 3681 | 2358 | 1952 | 802 | 898 | 2.8 | 3.9 | 0.27 | 0.23 | 2 | 10 | 173 | 22 |
| **57** | 66 | M | White | Non-Hispanic | 6 | 21 | 180 | 159 | 4 | 2 | N | 9108 | 4987 | 9199 | 2622 | 954 | 884 | 23 | 4.6 | 1.7 | 0.35 | 19 | 128 | 2049 | 45 |
| **71** | 45 | F | White | Non-Hispanic | 12 | 48 | 202 | 154 | 0 | 2 | Y | 5207 | 4559 | 1606 | 998 | 723 | 860 | 3.5 | 6.6 | 0.14 | 0.33 | 21 | 33 | 112 | 65 |
| **72** | 42 | M | White | Non-Hispanic | 16 | 35 | 188 | 153 | 1 | 2 | Y | 24822 | 10485 | 24034 | 2095 | 4887 | 2407 | N/A | N/A | N/A | N/A | N/A | N/A | 3138 | 81 |
| **82**** | 46 | M | N/A | Non-Hispanic | 0 | N/A | N/A | 163 | 0 | 1 | N | 6472 | 5187 | 2667 | 3094 | 1125 | 846 | N/A | N/A | N/A | N/A | N/A | N/A | 131 | 20 |
| **88** | 41 | M | White | Non-Hispanic | 7 | 23 | 180 | 157 | 1 | 1 | N | 8263 | 6730 | 1789 | 2276 | 1546 | 903 | 4.7 | 9.3 | 0.56 | 0.31 | 7 | 186 | 425 | 56 |
| **96** | 48 | F | White | Non-Hispanic | 9 | 30 | 194 | 164 | 0 | 1 | N | 24147 | 15675 | 3959 | 1498 | 1099 | 965 | N/A | N/A | N/A | N/A | N/A | N/A | 928 | 206 |
| **107** | 53 | F | White | Non-Hispanic | 10 | 29 | 202 | 173 | 0 | 2 | Y | 7967 | 6298 | 1560 | 1025 | 915 | 850 | 3.8 | 6.3 | 0.49 | 0.18 | 64 | 76 | 297 | 87 |
| **115** | 65 | F | White | Non-Hispanic | 20 | 41 | 188 | 147 | 0 | 2 | N | 26997 | 11600 | 19944 | 2081 | 991 | 890 | 63 | 22 | 2.9 | 0.27 | 116 | 157 | 1128 | 432 |
| **125** | 51 | F | White | Non-Hispanic | 10 | 26 | 168 | 142 | 0 | 1 | N | 4498 | 4271 | 2234 | 1361 | 684 | 807 | 1.8 | 1.1 | 1 | 0.19 | 4 | 2 | 127 | 10 |
| **131** | 39 | M | White | Non-Hispanic | 5 | 25 | 191 | 166 | 0 | 0 | N | 4285 | 3911 | 1318 | 943 | 1201 | 1166 | 0.27 | 0.93 | 0.35 | 0.26 | 1 | 3 | 50 | 14 |
| **134** | 27 | F | White | Non-Hispanic | 16 | 22 | 171 | 149 | 0 | 0 | N | 8884 | 6818 | 7472 | 2068 | 1057 | 982 | 4.1 | 3.7 | 2.1 | 0.37 | 15 | 6 | 2701 | 263 |
| **149** | 41 | M | White | Non-Hispanic | 17 | 28 | 173 | 145 | 1 | 2 | N | 6275 | 3875 | 1422 | 1073 | 1058 | 842 | 10 | 3.4 | 0.48 | 0.21 | 69 | 151 | 495 | 28 |
| **157** | 50 | M | White | Non-Hispanic | 10 | 32 | 179 | 147 | 0 | 1 | N | 11979 | 8751 | 11125 | 2370 | 1969 | 1374 | 15 | 7.9 | 5.4 | 0.71 | 67 | 89 | 742 | 190 |
| **173** | 47 | M | White | Non-Hispanic | 5 | 53 | 185 | 132 | 0 | 2 | N | 9127 | 5004 | 12194 | 1660 | 1162 | 979 | 2.5 | 2.8 | 7.3 | 0.59 | 149 | 143 | 647 | 176 |
| **190*** | 54 | F | White | Non-Hispanic | 18 | 63 | 190 | 127 | 0 | 4 | Y | 16156 | 10408 | 4567 | 1664 | 1207 | 1107 | 43 | 18 | 0.9 | 0.42 | 102 | 81 | 598 | 165 |
| **192*** | 47 | F | White | Non-Hispanic | 44 | 62 | 190 | 128 | 1 | 3 | Y | 13879 | 9000 | 5894 | 1525 | 1819 | 1598 | 30 | 22 | 0.68 | 0.38 | 106 | 145 | 608 | 409 |
| **195** | 24 | M | White | Non-Hispanic | 18 | 42 | 191 | 149 | 0 | 2 | N | 14242 | 7933 | 3954 | 2055 | 1227 | 978 | 22 | 6.9 | 1.5 | 0.24 | 15 | 31 | 1315 | 106 |
| **222** | 28 | M | Asian | Non-Hispanic | 11 | 37 | 173 | 136 | 1 | 2 | N | 14063 | 6930 | 1132 | 723 | 2841 | 1612 | 7.9 | 2.7 | 0.37 | 0.18 | 14 | 17 | 865 | 50 |
| **287** | 47 | M | White | Non-Hispanic | 11 | 23 | 165 | 142 | 0 | 1 | N | 7442 | 4357 | 2873 | 1211 | 910 | 928 | 9.3 | 3.9 | 0.77 | 0.41 | 15 | 15 | 240 | 38 |
| **310** | 34 | F | White | Non-Hispanic | 17 | 35 | 185 | 150 | 0 | 2 | Y | 26782 | 15634 | 1554 | 1023 | 1435 | 1083 | 47 | 15 | 0.27 | 0.31 | 51 | 137 | 485 | 153 |
| **314** | 46 | M | White | Non-Hispanic | 11 | 43 | 184 | 141 | 0 | 2 | Y | 12475 | 7247 | 2431 | 1273 | 854 | 811 | 38 | 14 | 0.89 | 0.2 | 131 | 198 | 667 | 297 |
| **393*** | 69 | M | White | Non-Hispanic | 23 | 54 | 187 | 133 | 0 | 5 | N | 8729 | 5150 | 13320 | 1974 | 1075 | 892 | 14 | 6.1 | 2.5 | 0.27 | 13 | 51 | 715 | 144 |
| **394** | 48 | F | Multiple | Hispanic | 7 | 67 | 200 | 133 | 2 | 2 | N | 22856 | 12823 | 6178 | 1909 | 1009 | 1131 | 96 | 35 | 1.1 | 0.34 | 59 | 69 | 1281 | 282 |
| **403*** | 52 | M | Asian | Non-Hispanic | 18 | 39 | 174 | 135 | 1 | 4 | Y | 24462 | 13614 | 4060 | 3187 | 2107 | 1164 | 170 | 29 | 0.7 | 0.13 | 29 | 41 | 3888 | 179 |
| **470** | 28 | F | White | Non-Hispanic | 17 | 51 | 173 | 122 | 0 | 2 | Y | 6054 | 4894 | 2315 | 1798 | 1003 | 1025 | 5.3 | 3.1 | 0.2 | 0.16 | 90 | 86 | 50 | 14 |
| **478** | 31 | M | White | Non-Hispanic | 16 | 52 | 172 | 120 | 0 | 1 | Y | 6600 | 4083 | 3238 | 1824 | 1264 | 1283 | 7.1 | 3.3 | 1.8 | 0.24 | 33 | 41 | 263 | 15 |
| **501*** | 32 | M | Asian | Non-Hispanic | 18 | 53 | 192 | 139 | 0 | 4 | Y | 22775 | 8667 | 5272 | 1242 | 1557 | 1098 | N/A | N/A | N/A | N/A | N/A | N/A | 719 | 125 |
| **506** | 46 | M | White | Non-Hispanic | 12 | 59 | 178 | 119 | 1 | 2 | Y | 3036 | 2595 | 1205 | 975 | 1338 | 1041 | N/A | N/A | N/A | N/A | N/A | N/A | 10 | 10 |
| **537** | 52 | M | White | Non-Hispanic | 15 | 45 | 178 | 133 | 2 | 2 | Y | 11285 | 6443 | 2448 | 1083 | 1245 | 1192 | 13 | 8.6 | 0.56 | 0.23 | 89 | 52 | 923 | 986 |
| **539*** | 73 | F | White | Non-Hispanic | 19 | 55 | 209 | 154 | 1 | 5 | Y | 20337 | 9568 | 7505 | 1386 | 1714 | 2124 | 68 | 41 | 1 | 0.65 | 144 | 199 | 488 | 50 |
| **632** | 38 | M | White | Non-Hispanic | 10 | 43 | 168 | 125 | 0 | 2 | Y | 16796 | 9152 | 1766 | 1548 | 2415 | 1833 | 31 | 12 | 0.36 | 0.25 | 141 | 153 | 572 | 161 |
| **633** | 39 | M | White | Non-Hispanic | 8 | 57 | 182 | 125 | 0 | 1 | N | 8759 | 5108 | 1436 | 1224 | 2019 | 1404 | 4.8 | 3.3 | 0.25 | 0.26 | 121 | 157 | 135 | 32 |
| **664*** | 45 | F | White | Non-Hispanic | 17 | 42 | 192 | 150 | 0 | 5 | Y | 12698 | 6927 | 3357 | 1420 | 1440 | 1395 | 11 | 4 | 0.43 | 0.21 | 61 | 26 | 384 | 37 |
| **674*** | 41 | M | White | Non-Hispanic | 17 | 57 | 182 | 125 | 0 | 4 | Y | 36682 | 21702 | 3061 | 1141 | 1320 | 1218 | 251 | 57 | 0.4 | 0.13 | 45 | 66 | 1619 | 298 |

Sx = symptoms
* = hospitalized, **=asymptomatic
§ = Arterial hypertension (HTN), obesity (OB), diabetes mellitus (DM), asthma (A), chronic obstructive pulmonary disease (COPD), coronary artery disease (CAD), cancer (CX)
¶ = WHO Ordinal Scale for Clinical Improvement, COVID-19 Trial Design Synopsis
† = Persistent fatigue, dyspnea, athletic deficit, or ≥ 3 other solicited symptoms beyond 6 weeks from Sx onset
Reported data are median (range) unless stated otherwise
